# ‘High-Density-SleepCleaner’: An open-source, semi-automatic artifact removal routine tailored to high-density sleep EEG

**DOI:** 10.1101/2022.08.09.503268

**Authors:** Sven Leach, Georgia Sousouri, Reto Huber

**Affiliations:** Child Development Center and Pediatric Sleep Disorders Center, University Children’s Hospital Zurich, University of Zurich, Zurich, Switzerland; Institute of Pharmacology & Toxicology, University of Zurich, Zurich, Switzerland; Department of Child and Adolescent Psychiatry and Psychotherapy, Psychiatric Hospital, University of Zurich, Zurich, Switzerland

**Keywords:** artifact removal, high-density electroencephalography, sleep, graphical user interface, outlier detection

## Abstract

**Background:** With up to 256 channels, high-density electroencephalography (hd-EEG) has become essential to the sleep research field. The vast amount of data resulting from this magnitude of channels in overnight EEG recordings complicates the removal of artifacts.

**New Method:** We present a new, semi-automatic artifact removal routine specifically designed for sleep hd-EEG recordings. By employing a graphical user interface (GUI), the user assesses epochs in regard to four sleep quality markers (SQMs). Based on their topography and underlying EEG signal, the user eventually removes artifactual values. To identify artifacts, the user is required to have basic knowledge of the typical (patho-)physiological EEG they are interested in, as well as artifactual EEG. The final output consists of a binary matrix (channels x epochs). Channels affected by artifacts can be restored in afflicted epochs using epoch-wise interpolation, a function included in the online repository.

**Results:** The routine was applied in 54 overnight sleep hd-EEG recordings. The proportion of bad epochs highly depends on the number of channels required to be artifact-free. Between 95% and 100% of bad epochs could be restored using epoch-wise interpolation. We furthermore present a detailed examination of two extreme cases (with few and many artifacts). For both nights, the topography and cyclic pattern of delta power look as expected after artifact removal.

**Comparison with Existing Methods:** Numerous artifact removal methods exist, yet their scope of application usually targets short wake EEG recordings. The proposed routine provides a transparent, practical, and efficient approach to identify artifacts in overnight sleep hd-EEG recordings.

**Conclusions:** This method reliably identifies artifacts simultaneously in all channels and epochs.

## 1. Introduction

Electroencephalography (EEG), a commonly used technique to measure electrical brain activity, has long been, and still is, the gold-standard method to differentiate vigilance states. Thus far, standard criteria for sleep staging by the American Academy of Sleep Medicine (AASM) are based on EEG characteristics (Berry et al., 2017). In the last decades, the emergence of sleep as a local and use-dependent phenomenon has met wide consensus and, thus, established the importance of high-density (hd-) EEG measurements. While alternative imaging techniques, such as magnetic resonance imaging (MRI) offer higher spatial resolution, their costs, sensitivity to movements, and noise make them impractical for sleep studies. The complementary increased temporal resolution of hd-EEG renders it an optimal technology for current and future applications. Ranging from 64 to 256 channels, both basic science and clinical applications have benefited from the high resolution of EEG recordings (Pisarenco et al., 2014). For example, sleep hd-EEG enables the observation of local increases of slow-wave activity (SWA; EEG spectral power in the delta frequency range, 0.5 – 4.5 Hz) in areas explicitly involved during pre-sleep learning (Huber et al., 2004). Furthermore, the vast amount of spatio-temporal characteristics that can be extracted from sleep hd-EEG, such as slow waves, sleep spindles, as well as cross-freq coupling, facilitates the study of brain function and development in healthy (Gorgoni et al., 2020) as well as in clinical populations. Therefore, hd-EEG has become especially important in the growing field of clinical diagnostics and therapeutics, especially for neuropsychiatric disorders such as depression and attention deficit disorder, as well as for stroke, traumatic brain injury, epilepsy, and specific applications in intensive care units (Popa et al., 2020; Freismuth & TaheriNejad, 2022). In epilepsy, for instance, hd-EEG is used to locate the epileptic focus of sleep-related epilepsy syndromes, usually in the context of pre-surgical screening (Avigdor et al., 2021). However, a great challenge accompanying the increased number of electrodes is the large amount of data. A critical processing step inherent to the majority of EEG analyses is the identification and removal of artifacts (i.e., unwanted components embedded within the EEG signal). This is a challenge in itself, as artifacts in the EEG may originate from various sources.

Typically, EEG artifacts can be classified into two broad categories: physiological (or internal, intrinsic) and non-physiological (or external, extrinsic). Intrinsic artifacts in sleep EEG originate from physiological functions such as eye and body movements, muscle tension, cardiac activity, or respiration. On the other hand, non-physiological artifacts are related to hardware and environmental conditions and may originate from external sources such as power line noise (50/60 Hz), electrode malfunction, electromagnetic interference, or environmental noise (Mumtaz et al., 2021).

In general, there are two approaches to handle artifacts in sleep hd-EEG. Whole-time segments (i.e., epochs of 20 s or 30 s) can be discarded and not taken into account for further analysis, a suitable approach for overnight sleep EEG as it comprises several hours of data. In this approach, epochs containing artifacts can be manually identified, allowing full control of the data. Artifactual epochs are commonly identified during sleep staging, yet only in channels used for sleep staging. Consecutive channel-wise manual artifact identification is impractical due to the large number of channels and does not allow for a global topographical comparison among them. Automated, unsupervised procedures have also been suggested (Coppieters’t Wallant et al., 2016; Saifutdinova et al., 2019), however, while convenient, they lack transparency in the process.

Alternatively, artifacts that occur at the same time as neuronal signals can be computationally distinguished from brain activity and removed such that clean underlying neuronal signals are preserved. Common procedures include adaptive filtering, regression-based methods, frequency decomposition methods (wavelet transform decomposition, empirical mode decomposition), and hybrid approaches. Especially blind source separation techniques are frequently applied, including independent component analysis (ICA), canonical correlation analysis (CCA), and principal component analysis (PCA) (Jiang et al., 2019; Bisht et al., 2020; Kotte & Dabbakuti, 2020).

For wake EEG, the most widely used technique is ICA as it performs well on prominent, physiological artifacts, such as eye movements, cardiac or muscular activity. While some studies have used ICA in overnight sleep EEG (e.g., Siclari et al., 2018), its application in sleep EEG is infrequent and thus no systematic assessment is possible at this time.

Consequently, a reliable approach that gives the user full control of the data while being practical and time-efficient is currently lacking for sleep hd-EEG recordings. We aimed to address this gap by introducing a new MATLAB-based, semi-automatic artifact removal routine freely available in an online GitHub repository (Hd-SleepCleaner). It works with epoched data and provides a graphical user interface (GUI) specifically designed for sleep hd-EEG recordings. The GUI allows for the simultaneous screening of all channels and epochs at once. The data is screened based on four different sleep-related signal quality markers (SQMs), reducing the workload significantly. It provides the functionality to visualize the EEG signal and topography of each SQM, offering full transparency of underlying EEG traces that can potentially be classified as artifacts. Taking all together, the proposed artifact removal routine constitutes an approach highly suitable for the cleaning of sleep hd-EEG recordings, by allowing the user full control over the removal process and new insights into their data.

## 2. Methods

### 2.1 Example data

The artifact removal routine (v1.0.0) was applied in 54 overnight sleep hd-EEG recordings (EGI Net Station v5.4; Electrical Geodesics Sensor Net for long-term monitoring, 128 channels, Net Amps 400 series, Electrical Geodesics Inc., EGI, Eugene, OR, USA). All channels were referenced to Cz during recording and sampled at a rate of 500 or 1000 Hz. EEG data was exported with a software specific 0.1 Hz high-pass filter (EGI Net Station v5.4) to remove slow drifts and voltage jumps.

EEG recordings came from 27 young and healthy good sleepers of average chronotype (6 male, 21 female, all right-handed, aged between 18.38 – 26.69 y, mean±sd = 22.68±2.23 y). Participants spent two nights in the laboratory, each offering an ∼8 h sleep opportunity. Neither they nor any family member had a history of neurological or psychiatric disease, including any sleep disorder. We demonstrate the functionality of the proposed artifact removal routine using two example recordings (20.23 y and 24.82 y, both female and right-handed). The data was collected within a study which investigated the effect of phase-targeted auditory stimulation on electroencephalographic parameters and cognition (Kantonale Ethikkommission Zürich, KEK-ZH, BASEC 2019-02134). The study was conducted in accordance with the declaration of Helsinki and written informed consent was obtained prior to participation.

### 2.2 Artifact removal routine

Each value in the summary plot corresponds to one epoch for a given channel. Consequently, outlier values correspond to an artifactual EEG signal from one channel in a given epoch. Outlier values are identified based on four sleep-related SQMs in the following order: 1) delta power (from robustly z-standardized EEG data), 2) beta power (from robustly z-standardized EEG data), 3) the maximum squared deviation in amplitude from the average EEG signal (from raw, not robustly z-standardized EEG data), and, 4) delta power (from raw, not robustly z-95 standardized EEG data). All four SQMs are computed from EEG data with original reference (EEG signal referenced as during recording), as well as from average referenced EEG data.

Hence, the artifact removal routine is repeated eight times. The first four times, the user iterates through all four SQMs from EEG data with original reference, thereafter through all four SQMs from average referenced EEG data. During each iteration, outlier values are identified and possible artifacts detected. As a final result, the artifact removal routine provides a matrix (channels x epochs) containing 0s and 1s, where 0 denotes that a certain channel contained artifacts or did not belong to the sleep stage of interest, and 1 indicates that a certain channel is artifact-free and within the sleep stage of interest. Thus, in total, each night is screened eight times for artifacts, each time based on a different SQM. The screening of one night takes approximately 10 to 60 minutes, depending on the length and quality of the data, as well as the experience of the user.

### 2.3 Requirements

#### 2.3.1. EEG requirements

For the artifact removal routine, the input EEG has to fulfill certain requirements. SQMs include spectral power in the delta (0.5 – 4.5 Hz) and beta (20 – 30 Hz) frequency range. In case the signal within those bands is not present or has been attenuated, those SQMs may not be sensitive towards outliers in the EEG. This is why the sampling rate, any filter, as well as the chosen epoch length need to meet certain criteria. The sampling rate needs to be high enough to reliably estimate beta power (minimum sampling rate of 60 Hz, Nyquist’s theorem). Any online (during recording) or offline (during analysis) filter should leave the delta and beta frequency band as unaffected as possible. The minimum epoch length is 4 s, as the routine computes spectral power in 4 s Hanning windows. Lastly, the artifact removal routine has only been tested in healthy sleep EEG. EEG from clinical populations, such as patients with epilepsy, may show a distinct pattern in chosen SQMs.

The artifact removal routine is specifically designed to screen sleep hd-EEG data consisting of 64 or more channels. While the routine still works with fewer and even one channel, the comparison of SQM values between channels will be more informative, the more channels are recorded. Note that the difference in amplitude from the channel average is meaningless when the data provided consists of only one channel.

The routine takes advantage of the naturally occurring time course of delta and beta power during sleep. In case EEG recordings of shorter length than one sleep cycle (e.g. naps) are screened for artifacts, special care is needed to investigate the EEG signal itself. The user needs to have a rather good understanding of artifactual and (patho-)physiological EEG data to be capable of distinguishing the two.

#### 2.3.2. Data format

Currently, the EEG needs to be stored in a .mat file, storing a common EEGLAB structure with EEG.srate and EEG.data as fields that contain the sampling rate and EEG data, respectively. The latter stores the EEG signal as a matrix (channels x samples). The function *makeEEG()* converges EEG data into an EEGLAB structure and is included in the online repository. Sleep stages need to be stored in a vector of numbers or letters, where a distinct number or letter corresponds to a certain sleep stage. Supported data formats currently include .mat, .txt and .vis files. A short example dataset (64 channels, one sleep cycle) is included in the online repository.

#### 2.3.3. System requirements

The artifact removal routine, as well as the GUI, were programmed using Matlab (R2021b, The Mathworks, Inc., Natick, Massachusetts) leveraging functions from EEGLAB (v2021.1; Delorme & Makeig, 2004). It is highly recommended using the same or a newer Matlab and EEGLAB version. Hardware requirements, e.g., RAM size, mainly depend on the size of EEG data that is processed.

### 2.4 Epoch selection

The artifact removal routine can selectively be performed only on epochs that belong to certain sleep stages (e.g. only NREM or REM sleep). This can be useful when only certain sleep stages are considered for analyses, as it lowers the number of epochs to be screened. Additionally, the hypnogram is displayed aligned to SQM values, providing useful information about their course in time. Epochs not belonging to the selected sleep stages of interest are treated as if they contained artifacts in the final output matrix.

### 2.5 Epoch-wise average referencing

Average referenced EEG data is only computed following the initial identification and removal of artifacts in the originally referenced EEG data. This is important as the average is susceptible to outliers. More specifically, amplitude values of artifactual channels are set to NaN in respective epochs. Then, the channel average used for average referencing is computed individually for each epoch. Note that for each epoch a different amount of channels may contribute to the average, as different channels can be artifactual in each epoch.

### 2.6 EEG preprocessing

The artifact removal routine comes with a simple EEG preprocessing pipeline, preparing the EEG data to correctly compute all SQMs. The EEG is low-pass filtered (−6 dB cut-off = 39.86 Hz, filter order = 92 at 500 Hz), down-sampled to 125 Hz (adjustable), and high-pass filtered (−6 dB cut-off = 0.37 Hz, filter order = 1494 at 125 Hz), primarily for the visualization of EEG traces and the computation of the maximum squared deviation from the average EEG signal. Spectral power in the delta and beta range should stay unaffected by the applied filters, as their passbands include the complete delta (from 0.5 Hz) and beta range (up to 30 Hz). When low-frequency, high-amplitude sweat artifacts contaminate the signal, stricter high-pass filtering (−6 dB cut-off = 0.75 Hz, filter order = 1246 at 125 Hz) is recommended for their removal. Sweat artifacts are high-amplitude, low-frequency waves which can lead to the exclusion of a significant amount of epochs during the artifact removal procedure. All filters are Kaiser-window-based FIR filters and are applied one-way with zero-phase shift. The stated filter order (in samples) is bound to the sampling rate of the EEG, so that doubling the sampling rate of the EEG would require doubling the filter order. The user has the option to directly load distinctively preprocessed EEG data into the routine and to skip the preprocessing step — a reasonable adjustment, as different filters introduce different filter artifacts (de Cheveigné & Nelken, 2019).

### 2.7 Normalization of EEG signals

The EEG measures the voltage difference between an active and a reference electrode. This is why the distance of the active to the reference electrode can impact the amplitude of the recorded signal considerably, much more than underlying brain activity. This is a problem when comparing amplitude-based SQM values among channels, as electrodes close to the reference will naturally have smaller amplitude values than electrodes further away (Fig. 3). Concomitantly, artifacts in electrodes close to the reference may be smaller than the physiological signal of electrodes further away from the reference electrode, encumbering their detection.

**Fig. 1.**
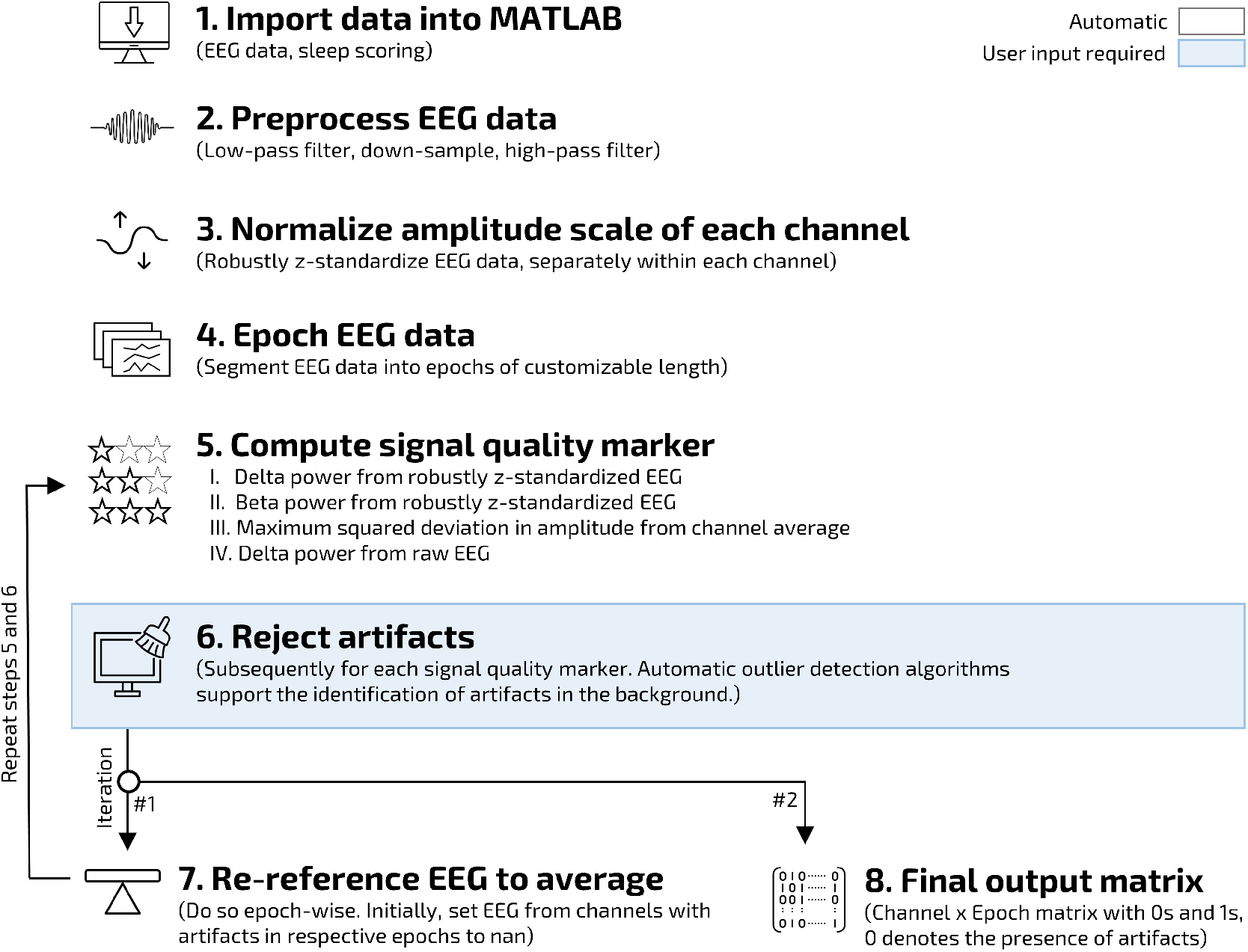
A flowchart depicting the artifact removal routine. After importing the EEG and sleep staging (optional), the EEG data is low-pass filtered, down-sampled, and high-pass filtered (optional) to correctly compute signal quality marker (SQM). To better compare SQM values among channels, the amplitude scale of each channel is normalized by robustly z-standardizing the EEG signal (channel-wise). For each epoch — usually of the same length as during sleep scoring — four SQMs are computed, three based on spectral power and one on the maximum squared deviation in amplitude from the channel average signal. Based on each of them, the user identifies conspicuous values, evaluates their EEG and topography, and, thereupon, removes values contaminated with artifacts. The EEG segments corresponding to rejected epochs for a given channel are set to NaN. Thereafter, the EEG is average referenced (epoch-wise), SQMs are computed once again, and values containing artifacts are removed. The final output matrix (channels x epochs) consists of 0s and 1s, where 0 denotes the presence of artifacts and 1 indicates clean EEG data.

**Fig. 2.**
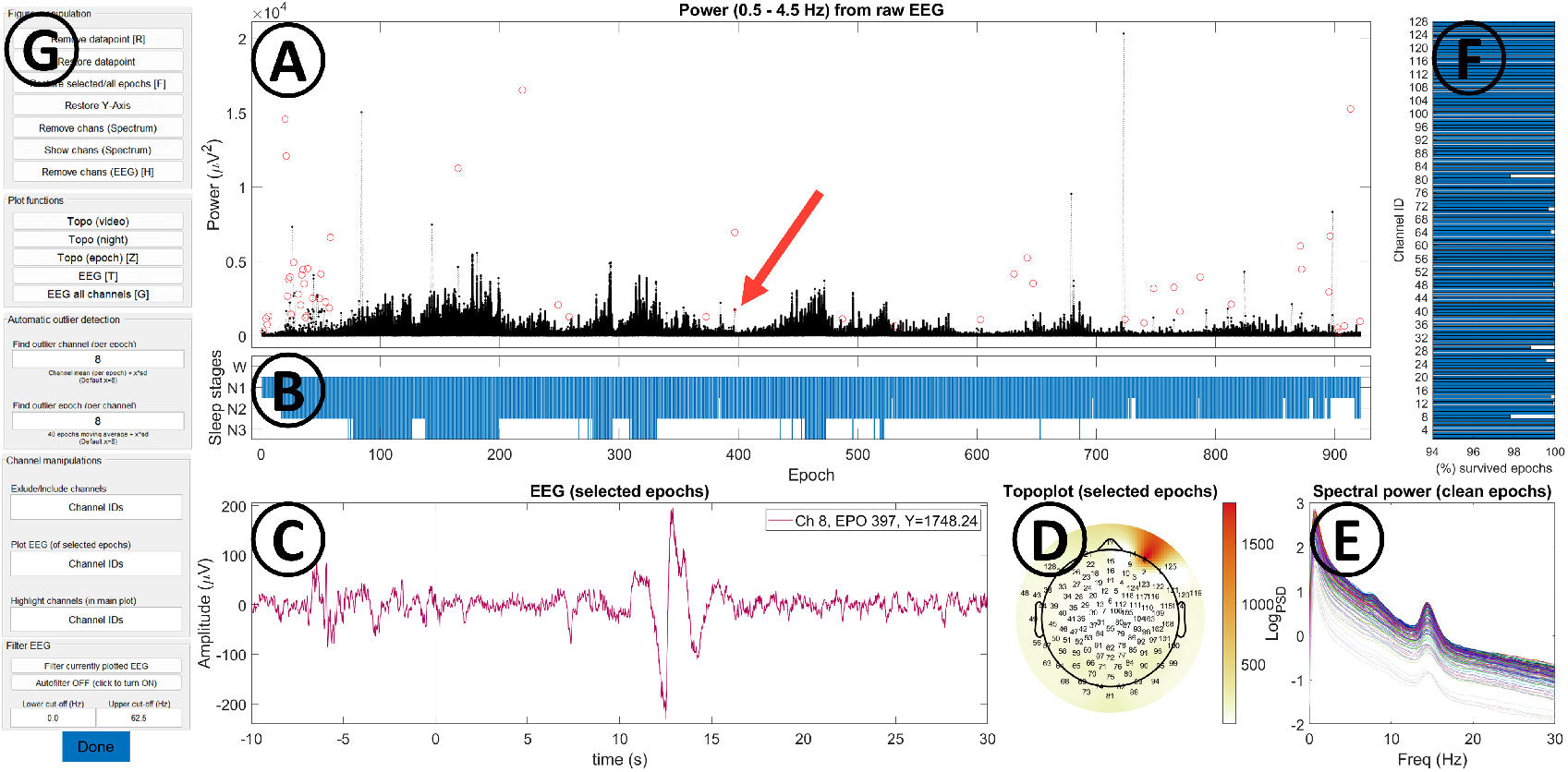
The graphical user interface (GUI). **A)** Each black dot represents a signal quality marker (SQM) value in one epoch for a given channel. SQMs include delta power (0.5 – 4.5 Hz), beta power (20 – 30 Hz) and the maximum squared deviation in amplitude from the channel average. Using the brush functionality (activated by clicking on Matlab’s specific brush icon), values can be selected and thereafter removed, visualized or restored. Removed data points are indicated by red circles. When pressing [DONE], each black dot is considered clean, each red circle artifact-contaminated. The red arrow in (A) indicates from which epoch the EEG in (C) and topography of SQM values in (D) is shown. **B)** Displaying corresponding sleep stages can help to assess the time course of SQM values. In case no sleep scoring is imported, no hypnogram is shown. **C)** The corresponding EEG signal of a selected SQM value (red arrow in (A) pointing to black dot). Evaluating the EEG signal is essential to determine the presence of artifacts. Artifactual EEG traces can be selected and removed. **D)** The topography of selected epochs can help to determine whether the source of SQM values is physiological or artifactual. Black dots in the topoplot indicate the corresponding channels of selected SQM values. **E)** Average overnight spectral power up to 30 Hz in steps of 0.25 Hz provides useful information about general channel quality. During NREM sleep, a peak in the delta and sigma band (12 – 16 Hz) are typically visible in all channels. The power spectrum automatically updates once SQM values are removed. **F)** The proportion of epochs which have survived the outlier detection routine for a given channel. Large amounts of removed epochs can indicate poor signal quality. **G)** Buttons allow interacting with the GUI. Letters in squared brackets correspond to keyboard shortcuts. The functionality of buttons is described in table 1.

**Fig. 3.**
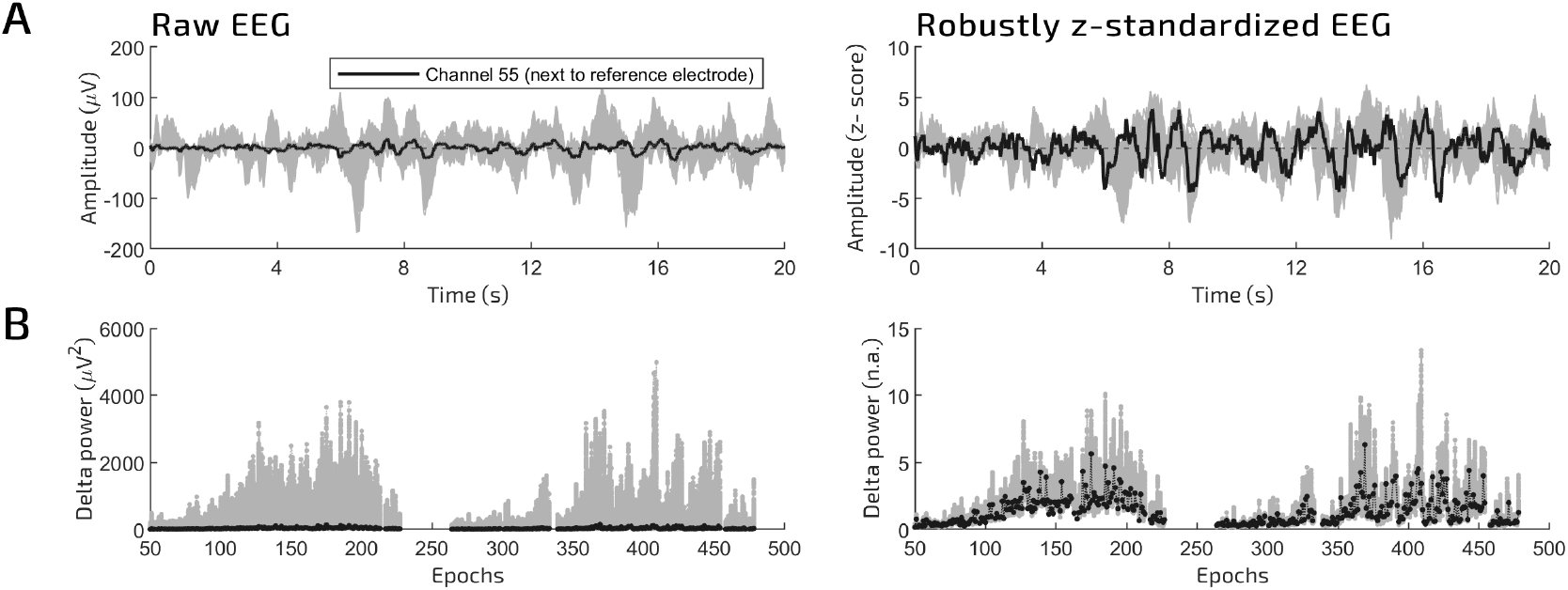
Robust z-standardization of each channel normalizes the EEG signal. **A)** (Left) Raw EEG signal (20 s) of all channels (gray). Channel 55 is highlighted in black. This channel is located next to the reference electrode and therefore exhibits relatively small amplitude values. (Right) After robust z-standardization of each channel, the amplitude of channel 55 during the same 20 s of EEG becomes larger relative to other channels. **B)** (Left) Delta power (0.5 – 4.5 Hz) of all channels (gray) during the first two sleep cycles. Channel 55 is highlighted in black and continuously shows one of the lowest delta power values compared to other channels. Smaller outliers are difficult to spot. (Right) After robust z-standardization of each channel, delta power of channel 55 is low in some and high in other epochs. This is due to the adjusted amplitude scale. Hence, robust z-standardization accounts for lower amplitude values for channels that are closer to the reference electrode during recording. The time course of delta power during the night (high in the beginning, low at the end of the night) does not change (not shown).

This is why it is essential to adjust the amplitude scale across channels when comparing channels altogether. Robust z-standardization over samples adjusts the amplitude scale of each channel individually 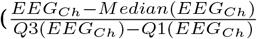, where *EEG*_*Ch*_ represents the whole-time signal of one channel). It uses the median and interquartile range (IQR) of the whole-time signal instead of the mean and standard deviation and is therefore robust towards extreme values which are naturally present in EEG data. A traditional z-score is the number of standard deviations by which the value of a raw score is above or below the mean value. A robust z-score can be interpreted analogously as the number of IQR by which the value of a raw score is above or below the median.

### 2.8 Spectral power computation

Three out of four SQMs are based on spectral power. For two of them, the EEG is robustly z-standardized over samples (separately for each channel). Spectral power is then computed for the delta (0.5 – 4.5 Hz) and beta (20 – 30 Hz) frequency range, a procedure used before (Huber et al., 2000) to capture artifacts well known to be present in sleep EEG, such as muscle tension, as well as eye or body movements. Delta power is computed both from robustly z-standardized and raw EEG data, the latter preserving the typical topography of delta power with a frontocentral hotspot during NREM sleep. Power spectral density (PSD) values, from which spectral power is calculated, are computed in each epoch with the Welch method using the *pwelch()* function in Matlab (4 s Hanning windows, 50% overlap, frequency resolution 0.25 Hz).

### 2.9 Maximum squared deviation in amplitude

The fourth SQM is computed in each epoch by deriving the maximum squared deviation in amplitude of one channel from the average EEG signal over channels (*max*(|*EEG*_*Ch*_ − *Mean*(*EEG*)|^2^), where *EEG*_*Ch*_ represents the signal of one channel for a given epoch and *EEG* is a data matrix (channels x samples) containing the EEG signal for a given epoch). Computing the squared amplitude amplifies differences and as such helps to detect smaller deviations. This measure is independent of the underlying frequency of artifacts, but instead detects deviations in amplitude of each channel from the channel average for a given epoch. Importantly, the attenuation of epoch edges, a necessary step for spectral power computations, is not required, making this SQM sensitive towards artifacts within an epoch just as well as at the edges.

### 2.10 Manual identification of artifacts

Eventually, SQM values are removed at each iteration whenever the user determines that their EEG signal is contaminated with artifacts. To confirm the presence of artifacts, it is crucial that the EEG and topography of outlier values can be visualized and examined, a feature the GUI provides. SQM values are indicative of artifactual sources when, 1) within the same epoch, a channel deviates from other channels, 2) within one channel, an epoch deviates from its neighboring epochs, 3) channels show a flat line (zero values), or, 4) the course in time of values over a night of sleep is not reasonable (e.g., higher delta power in the end than in the beginning of the night).

### 2.11 Visualizing the EEG

When selecting a conspicuous SQM value, the user has the option to 1) plot the corresponding EEG, or 2) plot the EEG of all channels from the same epoch. Alternatively, the user can select more than one SQM value and plot their corresponding EEG altogether. Visualizing the EEG is an essential attribute of the GUI. It contributes to confidently determine the presence of artifacts and makes the user aware of the specific types of artifacts present in their data.

### 2.12 Visualizing the topography of signal quality markers

The user can, furthermore, examine the topography of SQM values in a certain epoch, facilitating the identification of big deviations from neighboring channels specifically. When several epochs are selected at once, the maximum topography (maximum value over selected epochs) is displayed. Alternatively, the user can visualize several epochs successively in a movie format.

### 2.13 Automatic identification of artifacts

In the background and at the beginning of every iteration, two automatic outlier detection procedures support the artifact removal routine. More specifically, outliers are automatically detected 1) channel-wise, when values deviate more than X standard deviations from a moving average of 40 neighboring epochs and 2) epoch-wise, when values deviate more than X standard deviations from the average of all channels. Thresholds are adaptable within the GUI. In the presented data (128 channels, 8 h duration), they were situated between 6 and 12 standard deviations.

### 2.14 Additional GUI features

#### 2.14.1 Online filters

In some applications, it may be useful to assess whether artifacts could be removed by applying a filter. Slow waves, a high-amplitude, low-frequency signal, may be perfectly preserved even when high-frequency noise is present. The GUI therefore offers the option to individually low- and high-pass filter the plotted EEG signal. We implemented a minimum order Chebyshev Type II IIR filter, a steep and fast filter favorable for online applications, with adaptable pass- and stop-band frequencies. The user can select the stop-band frequency, the frequency from which (low-pass) or until which (high-pass) the signal is attenuated by at least 60 dB. The pass-band frequency, the frequency until which (low-pass) or from which (high-pass) the signal remains “unaffected”, is automatically set to 1.25 (high-pass) or 0.9 (low-pass) times the user-defined stop-band frequency. This approach helps to easily evaluate the EEG signal in the frequency range of interest.

#### 2.14.2. Average spectral power

The GUI shows the average log-transformed spectral power of all remaining epochs for frequencies between 0 and 30 Hz (0.25 Hz resolution) for all channels. Power spectra that deviate from other channels can be a hint of abnormal behavior in one channel. Normally, frontal channels show higher total spectral power compared to posterior channels. To remove this offset in power, the power spectrum is computed from robustly z-standardized EEG so that the power spectrum of all channels can be fairly compared. The user can select the power spectrum of one channel and then has the option to 1) highlight SQM values of that channel or 2) remove that channel completely. Whenever abnormal values are removed, the power spectrum updates automatically and is computed only from remaining epochs.

#### 2.14.3. Proportion of artifact-free epochs

The GUI provides the proportion of epochs (in %) that is labeled as artifact-free separately for each channel. This helps to assess to what extent a channel is affected by artifacts. Large amounts of removed epochs can indicate poor signal quality.

#### 2.14.4. Highlighting conspicuous channels

The user has the option to highlight one or more channels by providing the channel number. This is helpful to see the behavior of one specific channel in the course of the night. Abnormal SQM values in affected epochs can then be selected, visualized and, if necessary, removed as usual.

#### 2.14.5. Evaluation

The topography of SQM values before and after the removal of artifactual SQM values can be compared within each iteration. When doing so, the average topography of all (artifactual & artifact-free) SQM values is plotted next to the average topography of all artifact-free SQM values. Additionally, upon completion of the entire artifact removal procedure, a final output figure displays the time course of delta power, its average topography across the night, as well as the percentage of epochs which survived the artifact removal routine.

### 2.15 Further analyses

Eventually, the final output is a binary matrix (channels x epochs) with 0s (artifact) and 1s (clean), which can be independently used for further analyses. The four following options could be considered. A rather conservative approach is to only include epochs into the analysis in which all channels are artifact-free. This can lead, however, to the loss of a substantial amount of data, depending on its quality, which is usually tolerable for overnight EEG recordings, yet not always.

Alternatively, channels close to the neck can be ignored so that they do not contribute to the assessment of artifact-free epochs. This can save a good amount of epochs, as those channels usually contain more artifacts. Another option is to interpolate the complete signal of bad channels, which contain artifacts in over 3% (arbitrary number) of all epochs in sleep stages of interest. A combination of the two is also possible. Both lead, however, to the loss of clean data and do not help in case several channels across the scalp are responsible for the exclusion of epochs.

The option that restores the largest amount of epochs is epoch-wise interpolation. It interpolates channels and restores their data only in those epochs in which they were labeled as “bad”. In case more than two (arbitrary number) neighboring channels contain artifacts as well, the entire epoch is rejected, as interpolation is more meaningful when surrounding channels contain clean EEG. With this method, epochs in which only certain channels show artifacts can be included in the analysis. The online repository comes with the external function *Call_EpowiseInterp()* that performs epoch-wise interpolation when enough neighboring channels are artifact-free.

### 2.16 Application of the artifact removal routine in 54 overnight sleep hd-EEG recordings

The routine was applied in 54 overnight sleep hd-EEG recordings (128 channels). To assess the efficacy of the routine, the number of “bad” and “poor” channels, as well as “bad” NREM (N1 + N2 + N3) epochs was assessed.

#### 2.16.1. Classification of bad and poor channels

Channels were classified as “bad” when all corresponding NREM epochs contained unphysiological data. Consequently, bad channels do not exhibit a single artifact-free NREM epoch. Channels were classified as “poor” when less than 97% of all NREM epochs were artifact-free.

#### 2.16.2. Classification of bad epochs

The presence of a single artifact-contaminated channel within an epoch is sufficient to classify the epoch as “bad”. As a result, the proportion of bad epochs generally depends on the number of channels required to be artifact-free. The fewer channels required to be artifact-free, the higher the chances the epoch can survive.

Thus, for a holistic evaluation, bad epochs were classified based on two common subsets of channels, each requiring a different number of channels to be artifact-free. The stricter subset comprised 124 out of 128 channels and only excluded chin and cheek electrodes (excluded electrodes: E107, E113, E126, and E127, see Supplementary Fiig. 4). A common procedure to increase the number of artifact-free epochs is to exclude channels located in the outer ring of the hd-EEG net from further analyses. Channels of the outer ring are placed under the ear close to neck muscles and are therefore prone to artifacts. The more liberal subset included 111 channels and excluded channels located in the outer ring (excluded electrodes: E43, E48, E63, E68, E73, E81, E88, E94, E99, E107, E113, E119, E120, E125, E126, E127, and E128).

In addition, bad epochs were also classified when bad and poor channels were excluded from the respective subset. Poor channels can result in many, and a single bad channel in all NREM epochs to be classified as bad. The number of bad epochs is reported with and without poor and bad channels.

Bad epochs could result from a single, a few, or many bad channels present in the respective epoch. Therefore, for a given epoch, the amount of bad channels is quantified. Reported numbers are based on the stricter subset of channels (including bad and poor channels).

Eventually, bad epochs were interpolated using epoch-wise interpolation. The number of successfully interpolated epochs, as well as the final amount of artifact-free epochs, is reported. Bad epochs were classified based on the stricter subset of channels (including bad and poor channels).

#### 2.16.3. Survival rate of screened epochs

For a given channel, the survival rate of screened NREM epochs was computed. The survival rate was defined by the number of artifact-free NREM epochs over the number of all NREM epochs. The average survival rate of the stricter and more liberal subset of channels, as well as of channels located in the outer ring exclusively, is reported.

#### 2.16.4. Total number of artifacts

Eventually, the total number of detected artifacts was estimated. To this end, for all channels, the number of bad epochs was added up. Epochs in which all channels were classified as bad were not taken into account.

### 2.17 Epoch-wise interpolation in two example nights

Bad epochs were classified in a night with many and few artifacts, based on 124 out of 128 channels. Chin and cheek electrodes were excluded (E107, E113, E126, and E127). Bad and poor channels were included for the classification of bad epochs. For a given epoch, a single bad channel is sufficient to classify the epoch as bad. 120 out of 124 channels were allowed to be interpolated, excluding earlobe and mastoid electrodes (E49, E56, E57, and E100), as they intentionally capture less brain activity. Bad epochs were candidates for epochwise interpolation. Epochs with more than two neighboring bad channels were rejected. The number of artifact-free epochs before and after epoch-wise interpolation, as well as the number of rejected epochs, is reported.

## 3. Results

### 3.1 Robust z-standardization facilitates the comparison of signal quality marker values among channels

Signal amplitude was normalized by performing channel-wise, robust z-standardization on the EEG signal of each channel. The time course of the EEG signal stays unaffected by this procedure, as all values are multiplied and shifted by the same value. What changes instead is the amplitude scale. Before normalization, the amplitude of a channel close to the reference is close to zero, leading to minimal delta power values compared to other channels (Fig. 3). After normalization, the typical time course of delta power during a night of sleep is visible for all channels, independent of their distance to the reference electrode.

Even though robust z-standardization is vigorous towards outliers as it uses the median and IQR over the mean and standard deviation, very noisy channels may result in normalized values that can clearly deviate from other channels (usually smaller values). In other words, in case one electrode loses contact so that a substantial amount of data, e.g., 60% of the night is contaminated with high-amplitude noise, those high-amplitude values will lead to a higher median and IQR than usual. As larger median or IQR lead to smaller normalized values, normal-amplitude values will be smaller than intended (Supplementary Fiig. 1). To account for this scenario, the artifact removal routine additionally computes delta power based on raw (not robustly z-standardized) EEG.

Electrodes next to the reference electrode can also exhibit normalized values which deviate from other channels without artifacts being present. Because of the proximity to the reference electrode, very close electrodes record neuronal signals with low amplitude values. Hence, they are more vulnerable to technical and environmental changes, e.g., changes in impedance. Sometimes, this can lead to higher amplitude values after, e.g., the first sleep cycle, mimicking electrode malfunctioning after this point in time (Supplementary Fiig. 1). In this case, however, the electrode showed a physiological time course after average referencing (Supplementary Fiig. 2). As the signal is close to zero, average referencing technically overwrites the signal with the channel average, which represented a physiological time course. This highlights the importance of screening channels with different reference set-ups.

### 3.2 Conspicuous signal quality marker

The decision over which SQM values to remove is eventually made by the user. To this end, it is important to know which SQM values may result from artifacts. Generally, there are four possible scenarios of SQM values indicating artifactual EEG data (Fig. 4). (1) Artifacts that affect several channels, such as body movements, result in a pattern where SQM values from several channels deviate from their neighbors in one epoch (bad epoch). In case the artifact lasts longer than the duration of the epoch, more than one successive epoch can behave accordingly. (2) Artifacts that affect a single channel, such as electrode malfunction, result in SQM values from one channel which continuously deviate from all or most other channels over several epochs (temporarily bad channel). (3) In case one channel shows a flat line, e.g., due to loose contact or high impedance, the SQM value is zero (temporarily flat channel). (4) Many times, a single or a few channels contain artifacts in one or another epoch, e.g. due to temporary failures in the contact between the EEG sensor and the scalp. Those artifacts can be detected by spotting single outlier values (single outliers).

**Fig. 4.**
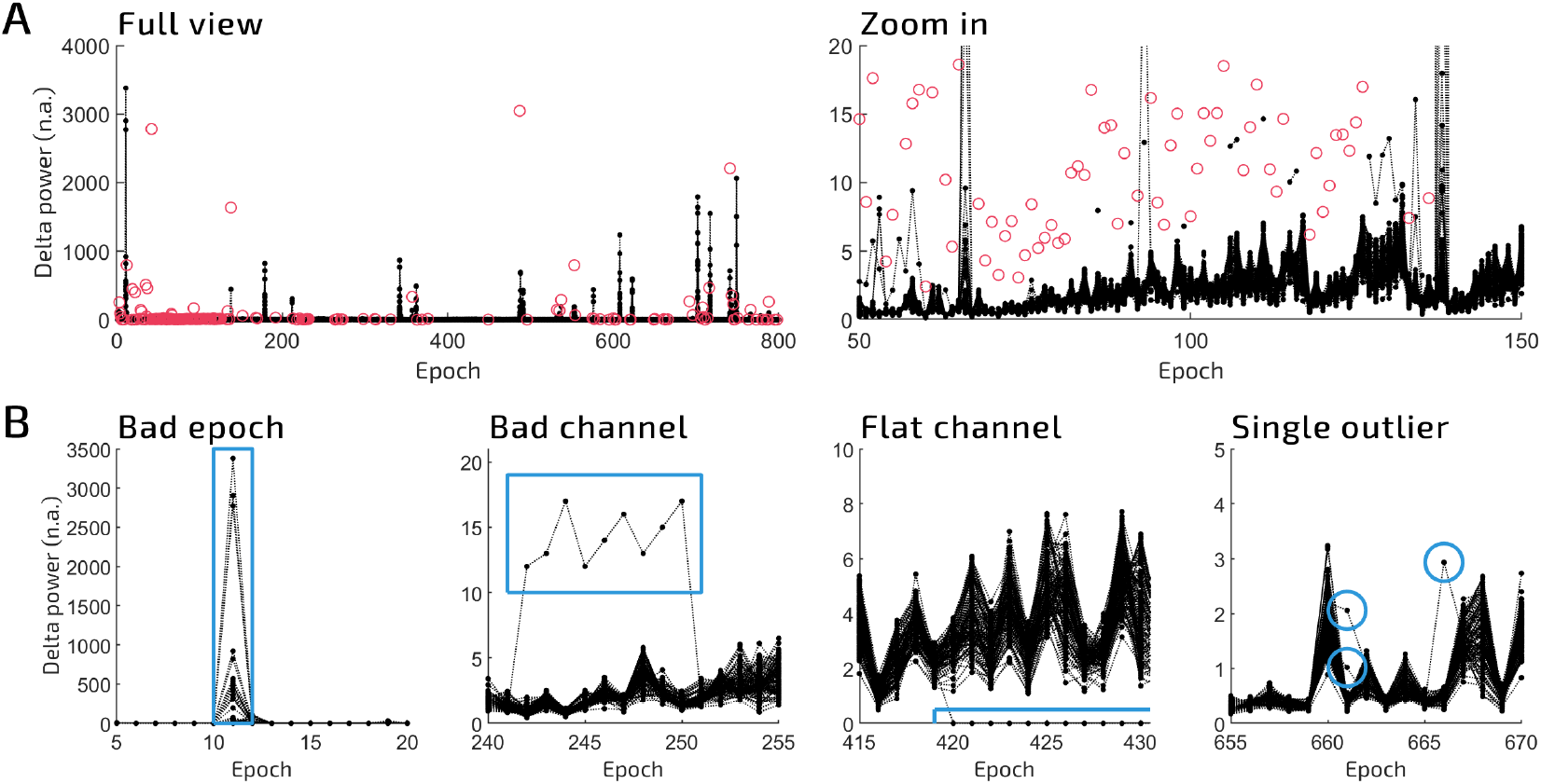
How to identify signal quality marker values with potential artifacts. **A)** (Left) Delta power (0.5 – 4.5 Hz) from robustly z-standardized EEG of all channel during all NREM (N1 + N2 + N3) epochs (20 s). One black dot represents the value in one epoch for a given channel. Red circles indicate values which were automatically removed by either of the two automatic detection algorithms, each worked with the default threshold of 8. (Right) Zoom into the first sleep cycle. Other conspicuous signal quality marker (SQM) values are now visible. **B)** Typical examples of SQM values which likely contain artifacts. Highlighted in blue, from left to right: 1) Compared to neighboring epochs, this epoch shows larger values in several channels, 2) one channel deviates from all other channels in several epochs, 3) one channel shows zero values in several epochs, 4) single values deviate from most other channels.

### 3.3 Example artifacts

To confidently determine the presence of artifacts, the user eventually evaluates the underlying EEG and topography of SQM values. We present five examples of common artifactual waveforms, as well as the corresponding topography of SQMs (Fig. 5). Even though the topography helps to localize and interpret the source of artifacts, we deliberately focus on the description of the waveform of the EEG, as the waveform is more relevant to determining the mere presence of artifacts. Low-frequency waves, including eye artifacts, are detected by deviations in delta power; high-amplitude noise of any frequency can be detected by the maximum squared deviation in amplitude from the channel average; high-frequency noise, as well as sharp waves, are detected by deviations in beta power. As the maximum squared deviation in amplitude from the channel average is independent of frequency, it can be sensitive to artifacts detected by delta and beta power, as well.

**Fig. 5.**
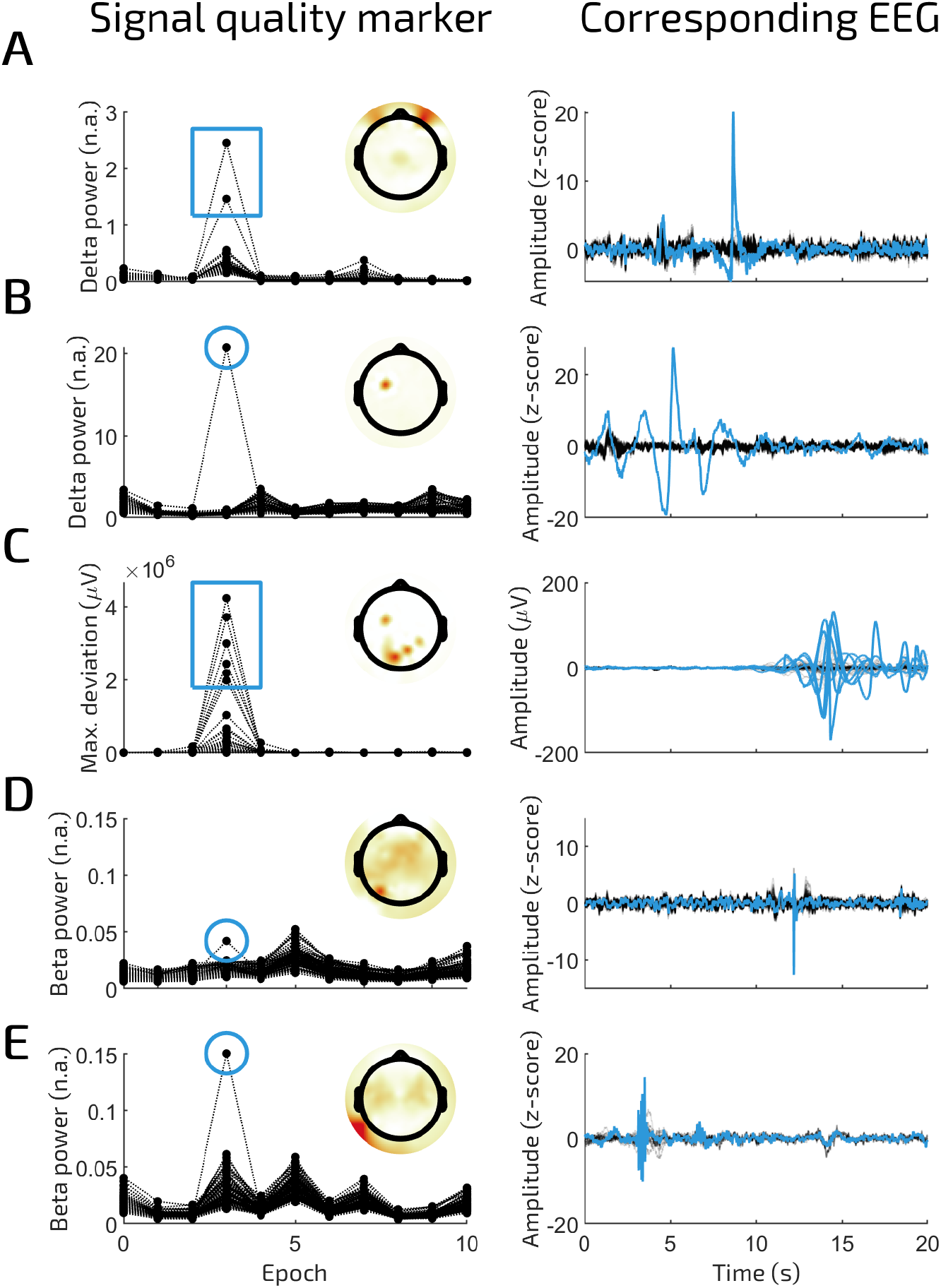
Example artifacts detected during the artifact rejection routine. **A)** (Left) Two prefrontal channels show high delta power (0.5 – 4.5 Hz) in one epoch (20 s). The topography points to eye artifacts. (Right) A single high-amplitude wave resembling an eye blink. **B)** (Left) A central channel shows high delta power in one epoch. (Right) The corresponding EEG shows slow, high-amplitude waves. **C)** (Left) Several posterior channels show high maximum squared deviation in amplitude from the channel average. (Right). The corresponding EEG shows high-amplitude noise. **D)** (Left) A posterior channel shows high beta power (20 – 30 Hz) in one epoch. (Right). The corresponding EEG shows a sharp, high-amplitude wave. **E)** (Left) A posterior channel shows high beta power in one epoch. (Right). The corresponding EEG shows high-frequency noise.

In some scenarios, however, the decision of whether the EEG signal is artifactual or physiological can be challenging. In sleep, a common scenario is when one or several channels show elevated beta activity for an extended period of time (Supplementary Fiig. 3). One possible reason is that participants lay on a specific electrode and thus apply pressure on the electrode. While the signal looks noisier than usual, frequencies of interest may be perfectly preserved (such as delta waves). Depending on further analyses, it may or may not make sense to label this data as artifactual, a decision that could lead to an increased number of excluded epochs. To give the user the possibility to assess how their frequencies of interest are affected, the GUI includes the functionality to filter the data in the desired frequency band of interest.

### 3.4 Evaluation of the artifact removal routine

To evaluate the success of the artifact removal procedure, we examined delta power, as well as log-transformed power spectral density (PSD) before and after the artifact removal routine. A hd-EEG sleep recording with many (bad night) or few artifacts (good night) was evaluated (Fig. 6). The bad night shows an unphysiological topography of delta power before artifact removal and immense outliers when looking at the time course. After the rejection of artifacts, both the topography and time course of delta power look as expected and physiological, with a fronto-central hotspot typical for young adults during NREM sleep after average referencing the EEG. After artifact rejection, delta power shows a common cyclic pattern which decreases over the course of the night, typical for sleep cycles during NREM sleep.

**Fig. 6.**
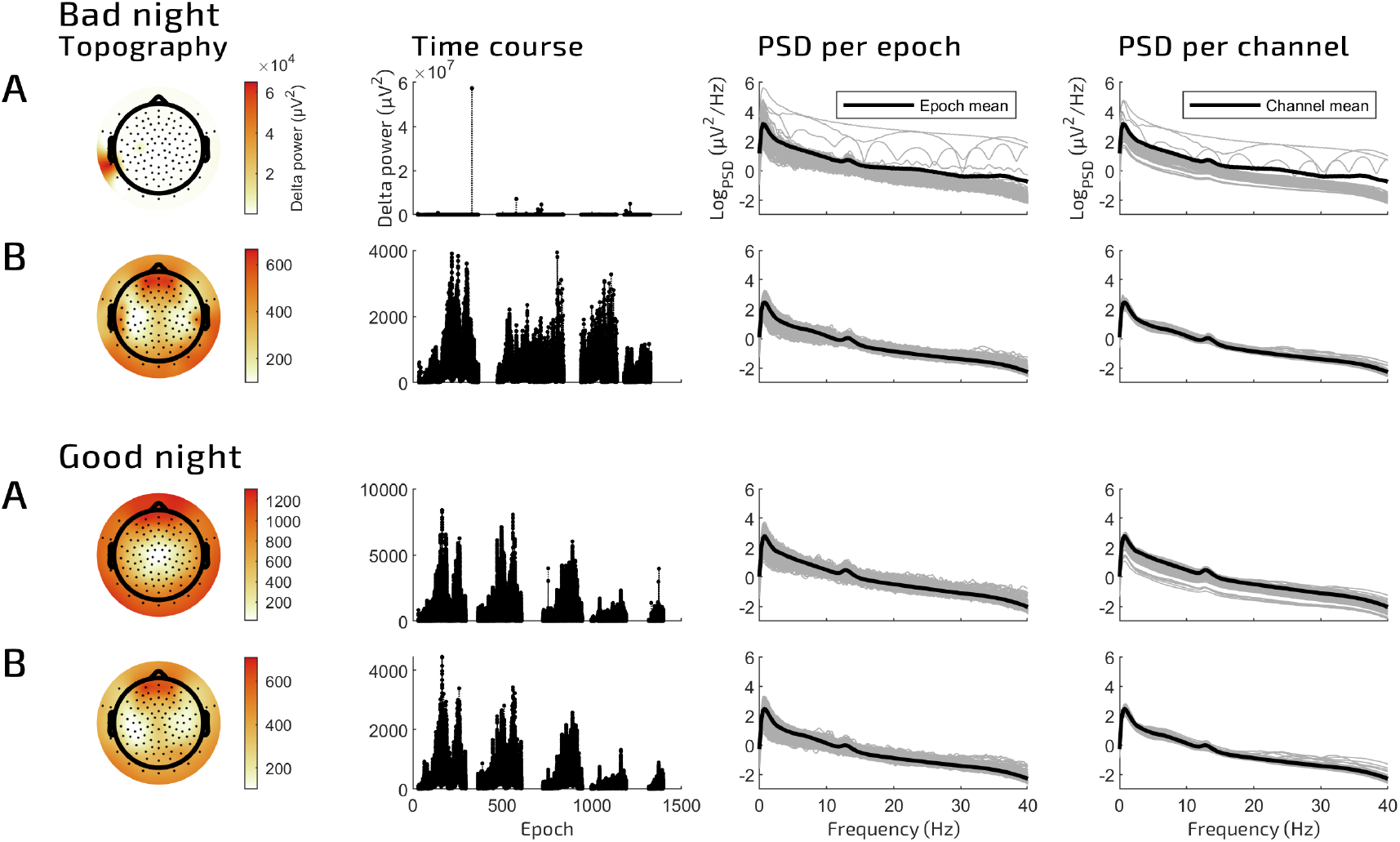
Spectral power before and after the artifact rejection routine. The figure shows spectral power of a sleep hd-EEG recording with many (top) and few artifacts (bottom). **A)** Spectral power before the artifact rejection routine. EEG data contains artifacts and is CZ referenced (as during recording). From left to right: 1) topography of average whole-night delta power (0.5 – 4.5 Hz), 2) time course of delta power during all NREM (N1 + N2 + N3) epochs. Each black dot represents the value of one channel in one epoch (20 s), 3) log-transformed power spectral density (averaged over channels, then log-transformed) of all NREM epochs between 0 and 40 Hz. One gray line represents one epoch, the black line the average of all epochs, 4) log-transformed power spectral density (averaged over epochs, then log-transformed) of all channels between 0 and 40 Hz. One gray line represents one channel, the black line the average of all channels. **B)** Same as (A) after the artifact rejection routine. EEG data is average referenced.

The same night shows several epochs with excessively high PSD (channel-average) for all frequencies between 0 and 40 Hz. After artifact removal, those epochs were removed and the variance between epochs reduced significantly. Moreover, a clear peak in the delta and sigma (12 – 16 Hz) range is visible. When comparing PSD among channels, some channels show excessively high PSD values (epoch-average). After artifact removal, all channels show similar PSD values with a physiological peak in the delta and sigma range.

In the good night, the topography and time course of delta power, as well as PSD look physiological and show very few outliers already before the artifact removal procedure. Only a few artifacts were removed, which is why delta power and PSD are similar before and after the artifact removal routine.

### 3.5 Application in 54 overnight sleep hd-EEG recordings

The proportion of bad epochs generally depends on the number of channels required to be artifact-free. Bad epochs were classified based on both a subset of channels that included, and a subset of channels that excluded the outer ring of the hd-EEG net. The subset including the outer ring resulted in more bad epochs (mean±sd = 46.77±35.07%, see Fig. 7A) than the subset excluding the outer ring (mean±sd = 31.91±26.86%). When bad channels were excluded, both the subset with (mean±sd = 25.79±16.15%) and without the outer ring (mean±sd = 23.93±15.86%) show approximately the same number of bad epochs. This indicates that bad channels are primarily located in the outer ring of the hd-EEG net. Indeed, more nights suffered from at least one bad channel when including the outer ring (*N* = 14 nights) than when the outer ring was excluded (*N* = 5 nights). Further excluding poor channels from both subsets of channels led to the lowest number of bad epochs, both for the subset with (mean±sd = 10.68±6.77%) and without the outer ring (mean±sd = 9.82±6.78%)

**Fig. 7.**
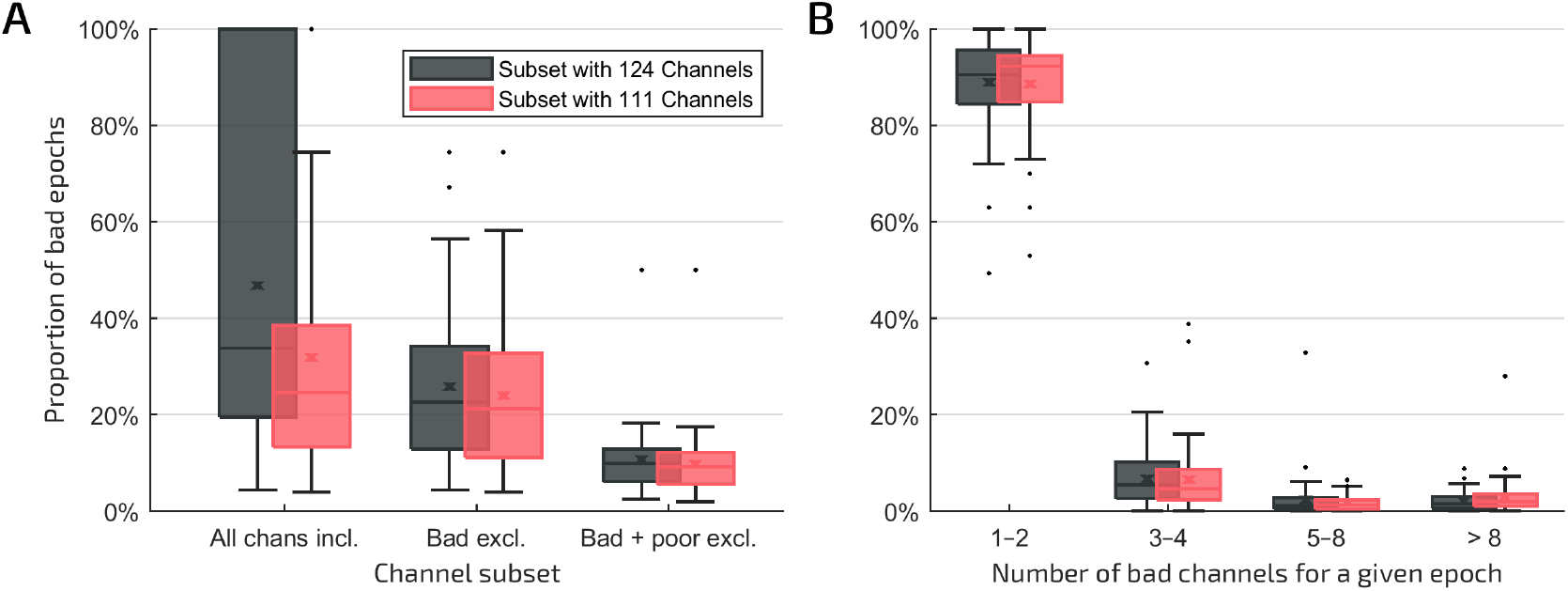
Proportion of bad epochs in 54 overnight sleep hd-EEG recordings. The artifact removal routine was applied in 54 overnight sleep hd-EEG recordings (128 channels). **A)** The proportion of classified bad epochs depends on the number of channels required to be artifact-free. A subset of 124 channels included for the classification of bad epochs leads to more classified bad epochs than a subset of 111 channels. Once bad channels are excluded, both subsets result in approximately the same proportion of bad epochs. The additional exclusion of poor channels further decreases the proportion of bad epochs in both subsets of channels. **B)** Most bad epochs only contain 1 – 2 bad channels. Bad epochs were classified with bad and poor channels included in both subsets of channels. *Boxplots:* The horizontal line indicates the median, the x the mean, the whiskers the scores within 1.5 times the interquartile range, and the box the middle 50% of scores.

Classified bad epochs differed in the amount of bad channels they exhibited (Fig. 7B), with either 1 – 2 bad channels (mean±sd = 88.83±9.15%), 3 – 4 bad channels (mean±sd = 6.73±5.84%), 5 – 8 bad channels (mean±sd = 2.31±4.62%), or ≥9 bad channels (mean±sd = 2.13±2.00%). As most bad epochs contained only few bad channels, the majority of bad epochs could be recovered using epoch-wise interpolation (mean±sd = 96.44±3.06%, range: 95% – 100%), resulting in almost all NREM epochs to be artifact free (mean±sd = 98.89±0.90%).

In total, a minimal amount of channels was classified as bad (mean±sd = 0.28±0.49 channels), and a moderate amount as poor (mean±sd = 5.52±16.68 channels). Most nights had no bad channel (*N* = 40 nights). The rest exhibited one (*N* = 13 nights) or two bad channels (*N* = 1 night). Poor channels were present in most nights (*N* = 48 nights), with 1 – 2 poor channels (*N* = 21 nights), 3 – 5 poor channels (*N* = 18 nights), 6 – 9 poor channels (*N* = 6 nights), or ≥10 poor channels (*N* = 3 nights). The average survival rate of epochs within a single channel was high (mean±sd = 98.92±0.88%) and comparable between channels located in the inner (mean±sd = 99.05±0.89%) and outer ring (mean±sd = 97.84±2.96%).

Using the routine in 54 nights, a total of ∼23,269 artifacts were detected.

### 3.6 Epoch-wise interpolation can recover nights with many artifacts

Artifactual EEG signals of respective channels can be interpolated in respective epochs. In a bad night, 0 epochs (0.00%) were artifact-free in a subset of 124 channels (Fig. 8). After epoch-wise interpolation, 893 epochs (95.00%) were restored. 47 epochs (5.00%) had to be rejected due to clusters of bad channels (more than two neighboring electrodes), potentially impairing the effectiveness of interpolation. In a good night, many more, namely 800 epochs (86.21%) were artifact-free, and a further 122 epochs (13.15%) were recovered using epochwise interpolation. 6 epochs (0.65%) had to be rejected.

**Fig. 8.**
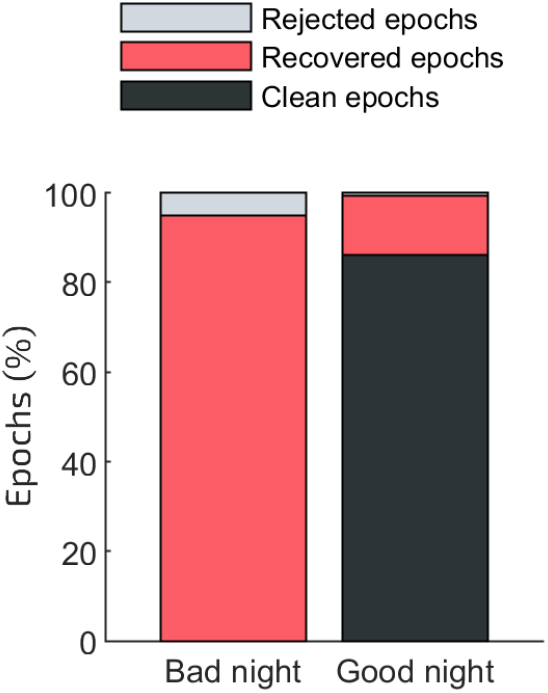
Proportion of rejected, recovered and clean epochs after epoch-wise interpolation. In an example night with many artifacts (bad night), no NREM (N1 + N2 + N3) epoch (20 s) was artifact-free in a subset of 124 channels. After epoch-wise interpolation, however, 95.00% of all NREM epochs could be recovered and used for further analyses. Without epoch-wise interpolation, those epochs had at least one channel containing artifacts. The other 5.00% of epochs were rejected. In an example night with few artifacts (good night), 86.21% of all NREM epochs were artifact-free in all channels and further 13.15% could be recovered using epoch-wise interpolation. A minimal proportion was rejected.

## 4. Discussion

Hd-EEG has become essential to the sleep research field and has started to enter sleep medicine, as well. This method has provided both basic and clinical science with the necessary tools to study local aspects of sleep, a large emerging area. Thus far, the resulting amount of data produced by recording several hours of sleep EEG with up to 256 channels still requests a practical and efficient approach to remove artifacts. In this work, we present a new, semiautomatic artifact removal routine specifically designed for sleep hd-EEG recordings, which allows the user full control of the data while being practical and time-efficient at the same time. Inside a GUI, the user assesses epochs based on four SQMs, evaluates their topography and underlying EEG signal, and, eventually, removes artifactual values. The possibility to visualize the actual EEG signal additionally familiarizes the user effortlessly with their data. The final output consists of a binary output matrix (channels x epochs) indicating artifactual and clean epochs with 0s and 1s, respectively. This output matrix can be used independently for further analyses of the data.

This artifact removal routine can be understood as a substantially advanced version of an established routine (first introduced in Huber et al., 2000) that has already been widely used in various sleep studies (e.g., Krugliakova et al., 2020; Sousouri et al., 2022; Lustenberger et al., 2015). The old routine screened channels and epochs based on delta and beta power (successively for each channel), an intuitive approach as artifacts typical for NREM sleep are in the delta (body movements, sweat artifacts, eye movements) or beta range (muscle activity, arousal). Artifactual epochs were removed by adjusting two thresholds, one defining an absolute power limit over which epochs were excluded, the other excluding epochs once absolute power exceeded a multiple of a sliding mean of an adjustable number of epochs. While this approach has proven efficient for the detection of evident outliers, it remains quite simplistic and less informative as compared to our multifaceted proposed method.

The presented routine also screens channels and epochs based on delta and beta power, yet additionally evaluates them based on the deviation in signal amplitude from the channel average, capturing artifacts independent of frequency, especially those present in single electrodes (i.e., loose contacts and other electrode malfunctions). As this SQM does not require the attenuation of epoch edges, it is sensitive towards artifacts at all locations within an epoch. Other major advances include the functionality to screen all channels at once, to manually remove artifactual values (without setting thresholds), to visualize underlying EEG traces, to evaluate the topography of SQMs, and to screen EEG data that is both Cz (original reference during recording) and average referenced. All these improvements aim to identify and remove as many artifacts as possible, while leaving physiological signal untouched. We successfully demonstrate that, when applying this artifact removal routine to both a night with many and a night with few artifacts, resulting average overnight delta power shows a physiological topography, epochs with artifactual delta activity are removed, and broadband spectral power shows a clear peak in the delta and spindle frequency range, typical for NREM sleep.

In total, the artifact removal routine was applied in 54 nights with varying data quality. An estimated 23,269 artifacts were found in 128 channels x 432 h of data. A minimal amount of channels was classified as bad by not containing a single artifact-free NREM epoch (0.28%). The few existing bad channels were usually located in the outer ring of the hd-EEG net. The survival rate of screened NREM epochs was high for single channels (98.92%). Yet, a single artifact-contaminated channel present within an epoch is sufficient to classify the epoch as bad. Accordingly, 9.82% to 46.77% of NREM epochs were classified as bad, depending on the subset of channels required to be artifact-free. The fewer channels were required to be artifact-free, the more epochs survived. Of those bad epochs, most exhibited only a minimal amount of bad channels. Hence, after epoch-wise interpolation, 98.89% of NREM epochs were clean, stressing the utility of this approach.

The wide range of classified bad epochs indicates that comparing the number of rejected NREM epochs to other artifact-removal techniques proves difficult. Not only does the number highly depend on the applied criteria (i.e., which channels are required to be artifact-free), but it also depends on the study population, number of recording electrodes, analyzed sleep stages, as well as data quality. It is important to note, however, that the routine encourages the user to keep as much physiological data as possible. Consequently, channels that exhibit artifacts for a considerable proportion of time are only excluded during artifactual time-windows. As a result, more epochs may be classified as bad at first, yet more physiological data can ultimately be recovered. This notably reduces the amount of data that needs to be interpolated. Other artifact removal approaches might reject the whole channel whenever a critical proportion of epochs is artifactual. While this may improve the number of artifact-free epochs as this channel will not be taken into account for the classification of bad epochs, it results in the loss of physiological data and is therefore less favorable.

This rigorous screening of artifacts demands the user to screen each night eight times (4 SQMs x 2 references). This is because some artifacts are only captured by one SQM specifically. Excluding one SQM from the routine would certainly lower the number of screening iterations, yet may also result in missed artifacts.

The artifact removal routine was designed to be as user-friendly as possible. To achieve this, two automatic outlier detectors shoulder the removal of extreme outliers. Moreover, the routine is completely modular with adjustable epoch length, optional sleep staging input, flexible sampling rate, and an optional preprocessing step included. Normalizing the EEG using robust z-standardization facilitates the evaluation of all channels altogether, which reduces the workload immensely while providing valuable insights for the detection of artifacts. Compared to powerful yet untransparent “black-box” machine learning approaches, which require training, this routine is completely transparent and only relies on optional thresholds.

Beyond the removal of artifacts, we provide a function that uses the final output matrix to interpolate artifactual channels in respective epochs. We show that this can save a great number of epochs, especially in recordings that suffer from bad data quality. By restricting the number of neighboring channels that can be interpolated, the function prevents the resulting loss of signal information to occur in a larger cluster of channels. It is important to note, however, that interpolation reduces the rank of the data (i.e., the number of independent signal sources). As a different number of channels is interpolated in each epoch, the rank of the data changes epoch by epoch, an important consideration for any further processing step that further relies on the number of independent signal sources. Furthermore, it is important to first remove voltage drifts by, e.g., high-pass filtering or detrending the data, before performing epoch-wise interpolation, as those voltage drifts might otherwise cause large edge artifacts.

### 4.1 Future directions

The online repository is constantly updated and improved. This includes the implementation of faster algorithms, novel functionalities, feature requests, and bug fixes. This process is expected to accelerate once more researchers use this routine for their own data. They may request additional features, raise issues, or even contribute their own code in the form of pull requests. The most current version is always accessible via the GitHub repository (Hd-SleepCleaner). This artifact removal routine is open-source and accessible to everyone.

Mid- and long-term plans for future directions include the exploration of a possible expansion to wake and REM hd-EEG recordings, as well as a possible integration of blind source separation techniques, such as ICA. ICA performs well at identifying and minimizing muscle (Crespo-Garcia et al., 2008) or ocular artifacts (Vigário, 1997), a type of artifact frequently present in REM and wake EEG recordings. Consequently, reducing muscle and ocular artifacts before performing the presented artifact removal routine could recover additional epochs.

### 4.2 Limitations

We have presented a simple, efficient, and effective artifact removal routine. Nevertheless, certain limitations are important to consider for each individual use case. This routine is semi-automatic and, therefore, provides full control over the removal process. However, it simultaneously requires the user to have at least some expertise in (patho-)physiological and artifactual EEG. As such, the final output matrix may vary depending on the user. The routine was specifically designed to screen all channels of hd-EEG recordings with at least 64 channels simultaneously. In turn, this routine may perform inferior with recordings including fewer channels for mainly three reasons: 1) the SQM which describes the maximum squared deviation in amplitude from the channel average is only meaningful when enough channels contribute to the channel average, 2) comparing SQM values of several channels within one epoch is more relevant when enough channels can be compared, and 3) the topography of SQM values is more informative the more channels contribute to the topography. Nevertheless, the functionality to visualize the underlying EEG of channels in respective epochs can prove valuable for recordings with fewer channels.

### 4.3 Availability

The artifact removal routine is open source and freely available on GitHub. The most current version is accessible via the online repository (github.com/HuberSleepLab/Hd-SleepCleaner) and website (HuberSleepLab.github.io/Hd-SleepCleaner). Please cite and refer to this paper whenever the artifact removal routine or parts of the online repository are used. The version used for analyses of this paper (v1.0.0) can always be traced back via the doi: 10.5281/zenodo.6883837. This work, as well as the online repository, run under the attribution license CC-BY. This license allows others to distribute, remix, adapt, and build upon the presented work, even commercially, as long as they credit this paper for the original creation.

### 4.4 Conclusion

The presented artifact removal routine allows the user to practically identify artifacts in overnight sleep hd-EEG recordings in a time-efficient manner for all channels simultaneously. After artifact removal, recordings show a topography and cyclic pattern of delta power as expected for NREM sleep. Up to 100% of epochs can be recovered using epoch-wise interpolation. The user is required to have basic knowledge of artifactual and (patho-)physiological EEG to differentiate the two.

## Acknowledgments

We thank the whole lab of Prof. Reto Huber for valuable feedback and discussions during the phase of development. Special thanks goes to Sophia Snipes for her exhaustive code review, suggestions for new features, and advice for the graphical user interface, as well as to Maria Eleni Dimitriades for her proofreading.

## Author contributions

SL developed the artifact removal routine, programmed the graphical user interface, recorded the overnight sleep hd-EEG recordings used for analyses, applied the routine and removed artifacts in 54 overnight sleep hd-EEG recordings, and performed all respective analyses. GS critically reviewed the source code and functionality of the artifact removal routine. SL wrote the paper together with GS. RH and GS provided oversight and leadership responsibility for the research activity, including mentorship to SL. RH critically reviewed the paper and acquired the financial support leading to this publication. This work was supported by the Swiss National Science Foundation (SNF, grant number 320030_179443).

## Data availability

Raw data that support the analyses of this paper cannot be made publicly available to protect participants’ rights according to Swiss human research law. The de-identified individual participant data that underlie the analyses of this paper can be accessed by investigators who (1) submit a methodological sound proposal describing the intended analysis and as reviewed by the authors of this publication, (2) provide proof of relevant ethical approval for the intended analysis, and (3) fulfill data protection measures according to Swiss legal requirements. The analyses of the shared data is restricted to achieve the aims of the intended analysis. Along with the data, no other documents will be made available. Proposals with reasonable requests may be submitted to the last author (email is not made public in the pre-print to avoid bot-spamming. Please search our names to get in contact or visit the github repository).

## ^2^Abbreviations

(SQM): signal quality marker
(hd-EEG): high-density electroencephalography
(GUI): graphical user interface
(NREM): non-rapid eye movement
(AASM): American Academy of Sleep Medicine
(PSD): power spectral density
(Q1): first quartile
(Q3): third quartile

## Tables

**Table 1:** Button functionalities. Several buttons are located on the left side of the graphical user interface (GUI), providing functionalities for the user to remove or restore signal quality marker (SQM) values, plot the EEG signal or topography of SQMs, adjust thresholds to automatically detect outlier values, highlight or remove whole channels, or to filter the plotted EEG signal in specified frequency ranges.

| *Buttons* | *Explanations* |
| --- | --- |
| Remove data points | Removes all selected values (black dots) within the main plot. |
| Restore data point | Restores one selected value (red circle) within the main plot. |
| Restore selected/all epochs | Restores all removed values (red circles) within the main plot. In case values (black dots) were selected, only restores those values that belong to the same epoch as the selected values. |
| Restore Y-Axis | Zooms out to restore Y-Axis. |
| Remove chans (Spectrum) | Removes all values (black dots) within the main plot from those channels whose "average overnight spectral power" was selected. |
| Show chans (Spectrum) | Highlights all values (black dots) within the main plot from those channels whose "average overnight spectral power" was selected. |
| Remove chans (EEG) | Removes all values (black dots) within the main plot from those channels whose EEG signal was selected. |
| Topo (video) | Displays the topography of values of all epochs subsequently, one after another, in a video-like format. In case values (black dots) were selected within the main plot, it only iterates through the corresponding epochs. |
| Topo (night) | Displays the topography (average over epochs) before (using all values) and after artifact rejection (using only artifact-free values). |
| Topo (epoch) | Displays the topography of the epoch a selected value belongs to. When several epochs are selected at once, the maximum topography (maximum value over selected epochs) is displayed. Black dots within the topoplot indicate which channels were selected. |
| EEG | Draws the EEG signal of selected values (black dots) within the main plot. |
| EEG all channels | Draws the EEG signal of all channels a selected value (black dot) belongs to. |
| Automatic outlier detection | Thresholds used for automatic outlier detection. Outliers are automatically detected 1) channel-wise, when values deviate more than X standard deviations from a moving average of 40 neighboring epochs and 2) epoch-wise, when values deviate more than X standard deviations from the average of all channels. |
| Exclude/include channels | Indicate channels. Removes all values (black dots) from indicated channels within the main plot. When all values of the indicated channels are already removed, it instead restores all values from those channels. In case one or more values (black dots) within the main plot are selected, it removes (or restores) only values of the same epoch the selected values belong to. |
| Plot EEG (of selected epochs) | Indicate channel indices and select values (black dots) within the main plot. Draws the EEG signal of indicated channels for those epochs the selected values belong to. |
| Highlight channels (in main plot) | Indicate channel indices. Highlights all values (black dots) of indicated channels within the main plot. |
| Filter currently plotted EEG | Applies a filter to the currently plotted EEG signal. |
| Autofilter ON (OFF) | Toggle button to automatically filter the plotted EEG signal. Can be turned ON or OFF. |
| DONE | Closes the GUI and continues the routine. |

## Additional material

### Graphical abstract

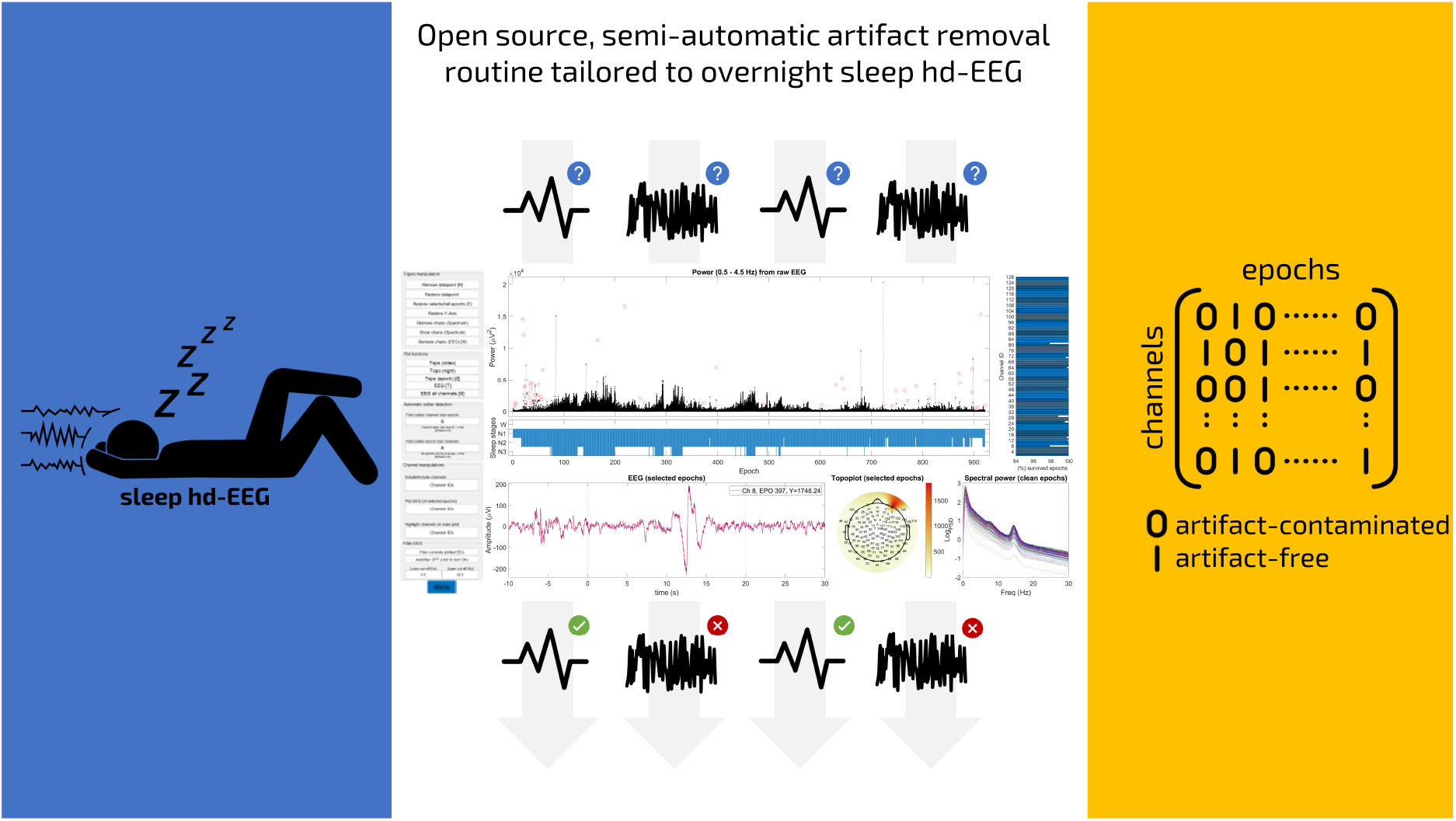

### Highlights

- Efficient semi-automatic artifact removal routine for sleep hd-EEG.
- Artifacts are identified in all channels and epochs inside a comprehensible GUI.
- The routine was applied in 54 recordings. Two example nights are assessed in detail.
- Clean epochs show a topography and time course of delta power typical for NREM sleep.
- Epoch-wise interpolation restores many epochs, especially when data quality is poor.

