## Supplementary material for "‘High-Density-SleepCleaner’: An open-source, semi-automatic artifact removal routine tailored to high-density sleep EEG"

### Supplemental information

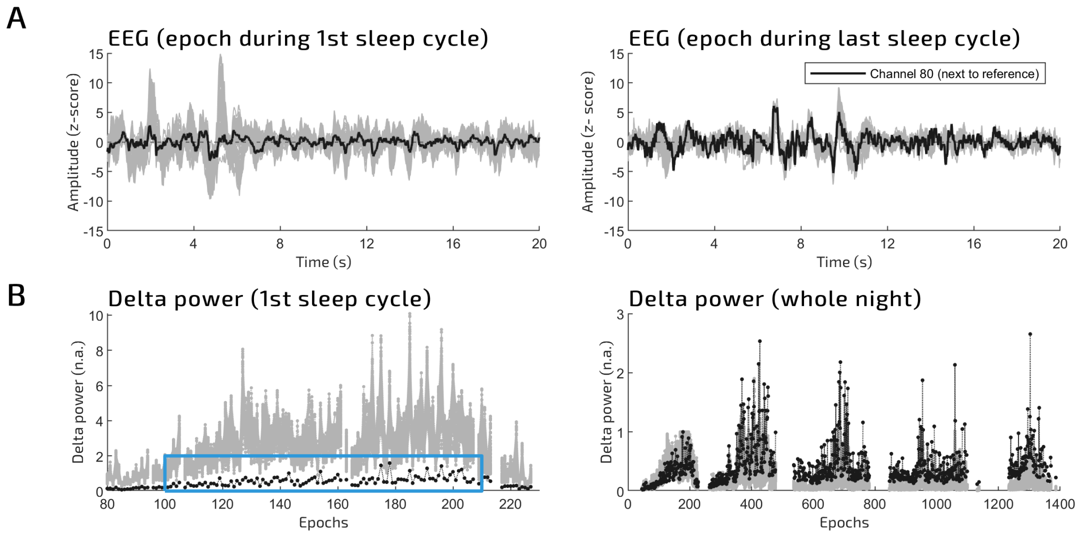


**Supplementary Fig. 1. Robust z-standardization is affected by amplitude changes. A)** (Left) Robustly z-standardized EEG signal (20 s) of all channels (grey) of an epoch during the first sleep cycle. Channel 80 is highlighted in black. (Right) Robustly z-standardized EEG signal (20 s) of an epoch during the last sleep cycle. After the first sleep cycle, channel 80 significantly increases in signal amplitude. **B)** (Left) Delta power (0.5 – 4.5 Hz) of all channels (grey) during the first sleep cycle. Channel 80 is highlighted in black and shows lower delta power values compared to other channels. (Right) When normalized to the maximum delta power value of the first sleep cycle, the time course of all channels becomes visible. For channel 80, delta power is lowest in the first and increases in later sleep cycles.

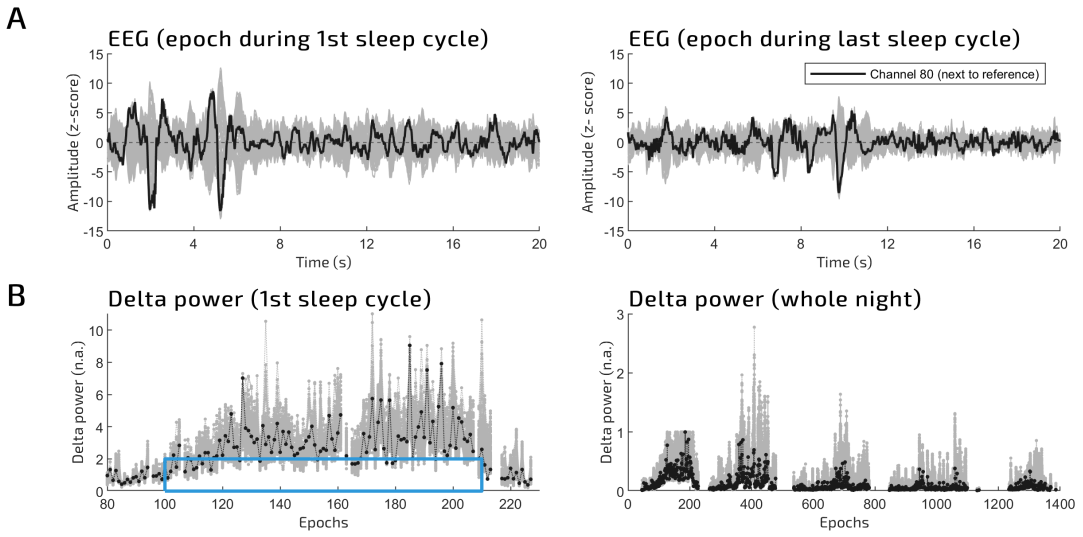


**Supplementary Fig. 2. Channels close to the reference electrode behave normally after average referencing. A)** Same as in Supplementary Fig. 1, yet the data was averaged referenced. Note that the amplitude of channel 80, a channel next to the reference electrode, behaves normally after average referencing. As the amplitude depends on the distance to the reference electrode during recording, the amplitude of channels next to the reference electrode is close to zero. When referencing to the average, their amplitude values are overwritten with the channel average **B)** Same as in Supplementary Fig. 1, yet note that delta power of channel 80 is now comparable to other channels after average referencing.

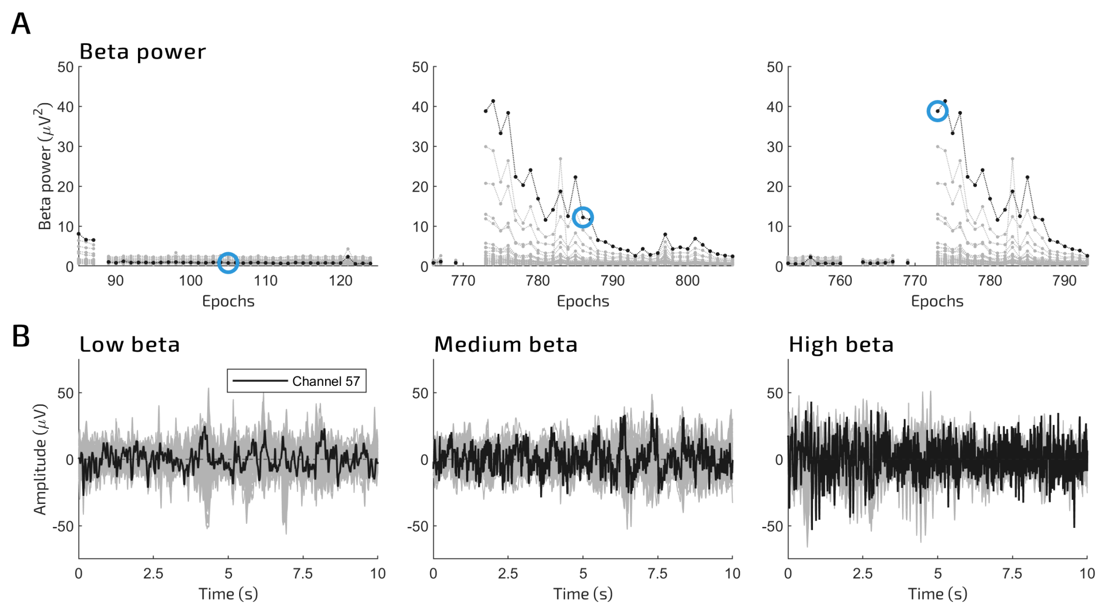


**Supplementary Fig. 3. Elevated beta activity.** Even though the channel clearly shows elevated beta activity, it can be difficult to judge whether the signal is artifactual and should therefore be excluded from the analysis.

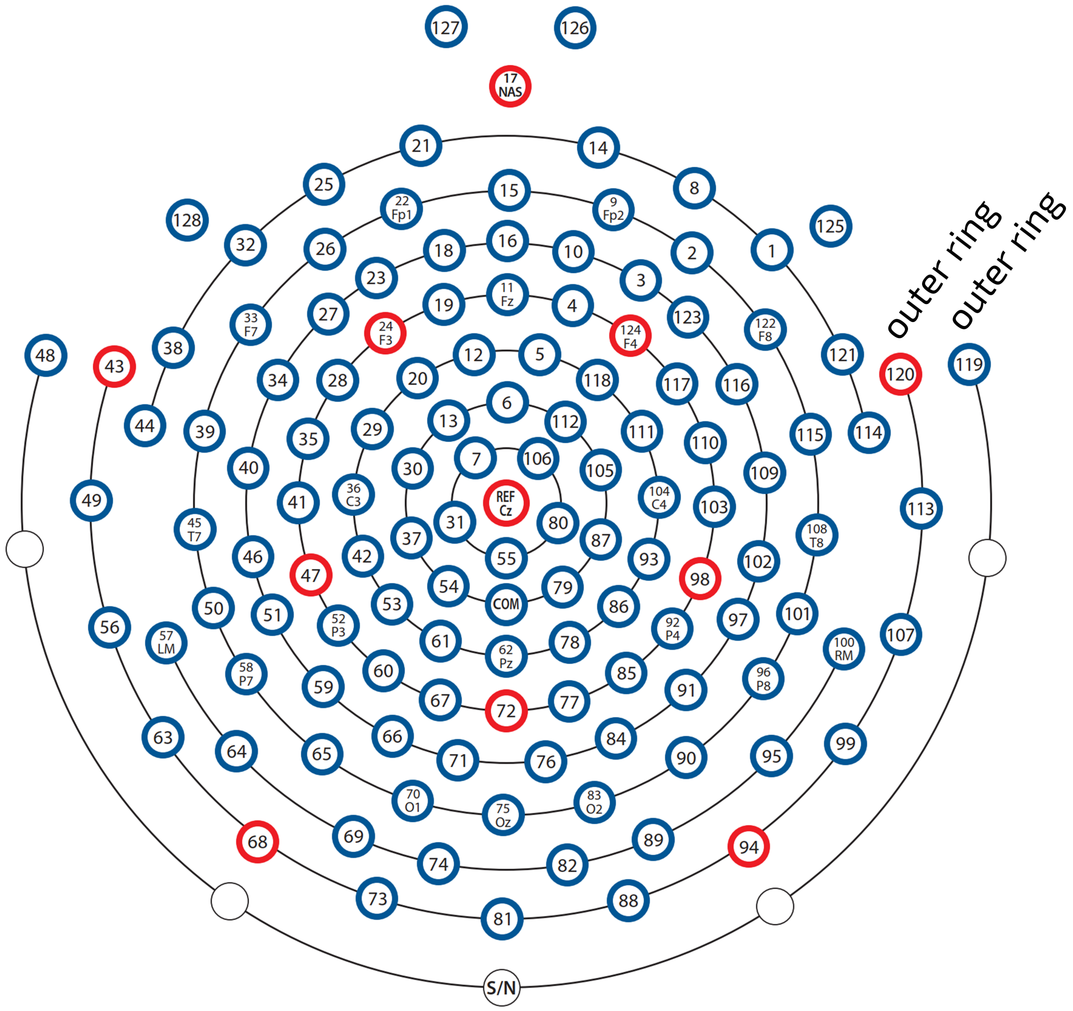


**Supplementary Fig. 4. Channel layout of the Electrocal Geodesics Sensor Net.** For some analyses, the outer ring of channels is excluded, corresponding to channels 43, 48, 63, 68, 73, 81, 88, 94, 99, 119, 120, 126, 127, 125, and 128. Note that channels 49 and 56, as well as 107 and 113 were external electrodes not integrated into the hd-EEG net. Instead, they were attached to the earlobes (49 & 56) and under the chin (107 & 113).
